# Assessing sex-specific selection against deleterious alleles: males don’t pay for sex

**DOI:** 10.1101/056663

**Authors:** Zofia M. Prokop, Monika A. Prus, Tomasz S. Gaczorek, Karolina Markot, Joanna K. Palka, Magdalena Mendrok

## Abstract

Selection acting on males can reduce mutation load of sexual relative to asexual populations, thus mitigating the two-fold cost of sex. This requires that it seeks and destroys the same mutations as selection acting on females, but with higher efficiency, which could happen due to sexual selection-a potent evolutionary force that in most systems predominantly affects males. We used replicate populations of red flour beetles *(Tribolium castaneum)* to study sex-specific selection against deleterious mutations introduced with ionizing radiation. Additionally, we employed a novel approach to quantify the relative contribution of sexual selection to the overall selection observed in males. The induced mutations were selected against in both sexes, with decreased sexual competitiveness contributing, on average, over 40% of the total decline in male fitness. However, we found no evidence for selection being stronger in males than in females; in fact, we observed a non-significant trend in the opposite direction. These results suggest that selection on males does not reduce mutation load below the level expected under the (hypothetical) scenario of asexual reproduction. Thus, we found no support for the hypothesis that sexual selection contributes to the evolutionary maintenance of sex.

## Introduction

The predominance of sexual reproduction among eukaryotic organisms remains one of the greatest puzzles in evolutionary biology: sex seems just too costly to be as common as it is. Sexual populations are typically composed of males and females, but only the latter invest resources directly into the production of offspring. Thus, an asexual (all-female) lineage will grow twice as fast as a sexual one if sexual females produce daughters and sons at 1:1 ratio and all else is equal between the lineages. This is the famous (or: infamous) two-fold cost of sex (Maynard Smith 1978, Milinski 2006), also called the cost of males.

However, all else is generally not equal: the presence of males can affect the reproductive output of females in a variety of ways. One intriguing theoretical scenario proposes that selection acting on males may mitigate or even eliminate the cost of sex (Manning 1984; Agrawal 2001; Siller 2001). Involved in that scenario is the concept of mutation load. Due to the incessant influx of mutations (Keightley and Lynch 2003), mean individual fitness in both sexual and asexual populations is lower than it would be for a mutation-free, Platonic ideal of a population: harmful alleles are maintained by the balance between the opposing forces of mutation and selection. This reduction in mean fitness is called mutation load (Agrawal 2013). Sex can push the mutation load of populations below the level experienced by the asexual ones if selection acting on males seeks and destroys (Hetfield et al. 1983) the same alleles as selection on females, and-crucially-does so more stringently (Manning 1984; Agrawal 2001; Siller 2001).

This could happen due to sexual selection, which arises from differential mating and fertilization success of individuals (Shuker 2010). Sexual selection is considered one of the most powerful and pervasive evolutionary forces (Andersson 1994; Kotiaho and Puurtinen 2007) and may often represent a major component of overall selection in sexual populations (Sharp and Agrawal 2012). Although it can (and does, in many taxa) act on both sexes, in most cases it tends to act much more strongly on males, limiting the number of those achieving paternity to the more competitive and/or attractive ones. Traits involved in male competitiveness and attractiveness are often energetically costly to produce; hence, their expression should depend on male condition (Price et al. 1993; Andersson 1994; Rowe and Houle 1996; Tomkins et al. 2004; Whitlock and Agrawal 2009). Condition of an individual can be defined as its overall health and vigour (Whitlock and Agrawal 2009) or more broadly, as a pool of resources it has acquired and can allocate into fitness-enhancing traits (Rowe and Houle 1996). As such, condition should be affected by a very large fraction of the genome, because almost every locus is likely to contribute, to some extent, to an organism’s ability to acquire and process resources. In that view, condition constitutes a “genetic exchanger” between male and female fitness, resulting in congruent direction of selection over most loci: most deleterious mutations will be deleterious precisely because they adversely affect condition, and in consequence-condition-dependent fitness components in both sexes (Rowe and Houle 1996). Crucially to the models’ (Manning 1984; Agrawal 2001; Siller 2001) assumptions, sexual selection on males could cause male fitness to be more sensitive to condition than female fitness, such that the fraction of reproducing males would be more limited (to those bearing relatively few condition-hampering mutations) than the fraction of reproducing females.

If these criteria are met, mutations detrimental to female fecundity are purged from sexual populations more efficiently than from asexual ones, “at the expense” of males (Agrawal 2001; Whitlock and Agrawal 2009). In effect, the cost of sex is counterbalanced by increased fitness of sexual females (Agrawal 2001). Importantly, this also improves the productivity of population, which is largely dependent on female fitness (Whitlock and Agrawal 2009). Alternatively, however, sexual selection may increase the unavoidable sexual antagonism, whereby some mutations favorable to one sex are disfavorable to the other (Connallon and Clark 2014). In effect, alleles detrimental to female fitness may be maintained in populations by selection acting on males (Plesnar-Bielak et al. 2014), further exacerbating the cost of sex (Holman and Kokko 2013).

Thus, sexual selection can have multiple effects on purging mutation load affecting female and population-level fitness-from lending “its aid to ordinary selection” (Darwin 1859) to counteracting it. Distinguishing between these scenarios is most frequently attempted by manipulating the intensity of sexual selection and analyzing downstream effects on the accumulation of the spontaneously arising mutation load (Radwan et al. 2004; McGuigan et al. 2011; Lumley et al. 2015) or the clearance of the experimentally induced one (e.g. Radwan 2004; Hollis and Houle 2011; Plesnar et al. 2011; Almbro and Simmons 2013; Power and Holman 2015). If sexual selection on males discriminates, by and large, against the same alleles as selection acting on females, then relaxing sexual selection should lead to a faster accumulation / slower clearance of mutations hampering female and population fitness, compared to treatments in which sexual selection is operating.

Importantly, however, such alignment is an insufficient-albeit necessary-condition for sexual selection to reduce mutation load in sexual relative to asexual populations, and thus contribute to paying the costs of sex. In order to chip in for the sex bill, sexual selection on males, at its naturaly occurring intensity, must act not only in the same direction, but also stronger, than selection acting on females (Agrawal 2001; Siller 2001; Whitlock and Agrawal 2009). This is perhaps most neatly explained in Whitlock and Agrawal’s (2009) review where the authors show that the expected mutation load (in a given locus) in sexual populations is as follows:

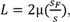

with μ being the rate of deleterious mutation, S_F_ - the coefficient of selection against deleterious mutations in females, and s-the average selection across the sexes, *i.e.* s = (S_M_ + S_F_) / 2; S_M_ being the coefficient of selection in males. In asexual populations, the predicted load is *L* = 2µ (Whitlock and Agrawal 2009). Thus, the sexual population’s load is reduced relative to the asexual case whenever the selection on females is weaker than the average selection across the sexes (which requires S_M_ > S_F_). A sexual population in which S_M_ = S_F_ is predicted to harbor the same level of load in female fitness as an otherwise identical asexual population (Agrawal 2001; Whitlock and Agrawal 2009). If the selection is weaker in males, the load is increased relative to the asexual case (see also a simple numerical example in the Supplement).

Thus, if the net effect of sexual selection on genome is in the same direction as of selection on females, relaxing the former will indeed hamper purging mutation load affecting female and population fitness, generating differences between treatments with relaxed vs. operating sexual selection, as observed in some studies (e.g. Radwan 2004; Lumley et al. 2015). However, such results cannot on their own verify that sexual selection does indeed improve fitness prospects of sexual females (and populations) compared to their asexual competitors. For this, it is necessary to determine not only the alignment of selection, but also its relative strength in males vs. females.

To date, this has been done by relatively few studies, and almost exclusively on fruit flies. For example, Sharp and Agrawal (2008) compared the strength of sexual selection against eight mutations with visible phenotypic effects in *Drosophila melanogaster*. They found significant sexual selection at low and high densities on six of eight mutations. They also found that for five of the examined mutations, selection on females was less important in eliminating them than selection on males. In another study, the same authors (Sharp and Agrawal 2012) measured sex-specific fitness effects of spontaneous mutations in *D. melanogaster,* using mutation accumulation lines. They found that accumulated mutations caused substantial fitness declines in both males and females with the effects being positively correlated between the sexes. Importantly, the mutations had larger effects on fitness in males than in females. Mallet et al. (2011), using X-chromosome mutation accumulation in the same species, also found stronger negative effects on males. Pischeda and Chippindale (2006) assessed the impact of deleterious *nub* mutation on *D. melanogaster* populations and found that mutant males experienced a greater decline in fitness than mutant females. Whitlock and Bourguet (2000) examined deleterious effects of five mutant alleles or pairs of alleles *(black, plexus-speck, claret, hairy, ebony-stripe)* with visible phenotypic effects on female productivity and male mating success. They showed that all except *hairy* had a deleterious effect on female productivity, whereas three (*black, claret, ebony-stripe*) were deleterious for male mating success; for those three, the effects on males tended to be stronger than on females. In a sole (to the best of our knowledge) example of a non-drosophilid study, Grieshop et al. (2016) induced mutations by gamma irradiation to compare sex-specific competitive lifetime reproductive success (LRS) in a seed beetle *Callosobruchus maculatus*. Male competitive LRS was strongly decreased by induced mutations whereas in females the effect was weaker and non-significant, indicating that selection against novel mutations was stronger on males than females.

Here, we analyzed the effects of mutations, induced by ionising radiation, on male and female fitness in the red flour beetle *Tribolium castaneum*. We induced the mutations on the genetic background of three replicate populations, which had been maintained at population size of 20 breeding adults for 46 generations prior to our experiment (as a part of a long-term project by another group, cf. Laskowski et al. 2015). They are therefore expected to harbor very little genetic variation, making it easier to detect the effects of the experimentally induced mutations (Pekkala et al. 2009). We induced mutations in adult males, and scored fitness effects in their offspring of both sexes. We only irradiated the fathers in order to minimize the influence of non-genetic trans-generational effects of irradiation. Thus, we analyzed the effects on male and female fitness of mutations inherited via their fathers’ germ line and thus present in the heterozygous state, mimicking the natural situation where mutations are rare (Radwan 2004). Our main aim was testing whether selection against deleterious mutations is indeed stronger in males than in females.

Additionally, we wanted to assess to what extent sexual selection itself contributed to the overall selection against deleterious mutations in males. We measured fitness as the number of adult offspring produced over one week of interactions with four individuals of the opposite sex and three competitors of the same sex. This type of design is commonly employed to assess fitness of males (and less frequently-females), although the numbers of individuals used in the assays vary among studies (Michalczyk et al. 2010, 2011; Lewis et al. 2012; Sharp and Agrawal 2012; Duffy et al. 2014). In males, the fitness outcome of such assay depends on their sexual competitiveness (the sexual selection component) and subsequent egg-to-adult offspring survival (offspring viability selection component). In our experiment, detrimental effects of mutations on male fitness could come about due either or both of these components. Thus, our second aim was to quantify the relative role of the sexual selection component. Finally, we conducted a behavioral assay to assess whether male sexual activity (one of key components of overall sexual competitiveness in flour beetles, (Michalczyk et al. 2011)) is affected by the induced mutations.

## Methods

### Introducing mutations

We induced mutations on the genetic background of three replicate populations (henceforth denoted as lines 50, 52, and 55), which had been maintained at population size of 20 breeding adults for 46 generations prior to our experiment, and were hence expected to harbor very little initial genetic variation (Laskowski et al. 2015). In each replicate, we applied three irradiation doses: 10, 20 and 40 Gy, plus a non-irradiated control (0 Gy). Three doses were used because we had not been sure *a priori* which would be sufficient to cause a detectable fitness decline in offspring.

Before the irradiation treatment, virgin males *(ca*. 7 days post eclosion) were individually placed in 6 cm Petri dishes filled with fodder, and mated to 4 females from a phenotypic marker (reindeer honey dipper, henceforth: RdHD) strain each. The RdHD strain is homozygous for a dominant allele causing exaggeration of antennal club, easily distinguishable by eye. These initial matings were introduced in order for the males to replenish their sperm reserves before the irradiation treatment, so that the offspring used for fitness assays were produced from germ line cells affected by irradiation. RdHD females were used so that they could be easily distinguished from the males, as the sexes are difficult to tell apart by eye in adult *T. castaneum*. After 7 days of interaction, the females were discarded, whereas the males were randomly assigned to treatment groups and (apart from the control group) irradiated with γ rays from a Cs-137 source at the Institute of Nuclear Physics (Polish Academy of Sciences) in Krakow.

Following the irradiation treatment, each male was individually placed in a 6 cm Petri dish filled with fodder, and mated to 2 virgin, non-irradiated females from the same line he originated from *(i.e*. line 50, 52 or 55). After 5 days of interactions, the males were discarded and each female was placed individually in a Petri dish with excess food, and left for a week to lay eggs. At pupal stage, the offspring of one randomly chosen female per each irradiated or non-irradiated male were isolated and separated by sex (two females per male were initially used only as a back-up in case one of them did not produce offspring). One son and one daughter per male were subsequently used in fitness assays.

### Fitness assays

As a proxy for fitness (W), we measured the number of adult offspring produced during one week in the context of a small population (design modified from the “reproductive success in population context” assay, (Lewis et al. 2012)) for one daughter and one son of each of the irradiated and non-irradiated males. The individuals used in these assays are henceforth referred to as mutated (offspring of irradiated males) and control (offspring of non-irradiated males) focal females and males, respectively.

Each focal male was placed in a container with 3 virgin males from the RdHD strain and 4 virgin females from the main (wild type) stock culture. Similarly, each focal female was placed in a container with 3 virgin females from the RdHD strain and 4 virgin males from the main (wild type) stock. These groups of beetles (henceforth referred to as experimental “populations”) were left to interact for 7 days. After that, we discarded the adults and added excess food to the containers for their developing offspring. We raised these offspring to adulthood, then killed them by freezing, separated by phenotype (wild-type or Rd) and counted. Since the RdHD allele is dominant and the RdHD strain we used is homozygous for it, we could unambiguously assign all wild type offspring the focal individual.

### Estimating selection against induced mutations in males and females

Initial data exploration revealed that 10 Gy dose did not produce any consistent decline in fitness of the irradiated males’ offspring of either sex, compared to controls (fig. S1). Therefore, we dropped this dose from further analyses.

In order to compare the strength of selection acting on males and females, we calculated the standardized coefficients of selection (s) against mutations induced by 20 and 40 Gy doses of γ rays, separately for each line and sex, as

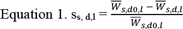

where the subscripts s, d and l denote sex, dose (with which fathers were irradiated; d0 stands for the control), and line, respectively, and 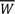 denotes mean fitness in a given group; e.g., 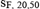 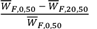. Then, for each dose and population we calculated the ratio of the s coefficients in males and females, m/f = S_M,d,l_ /S_F,d,l_ (Agrawal 2001).

In order to account for the uncertainty of the estimated coefficients, we used bootstrapping to obtain the confidence intervals around them (following the approach used by Sharp and Agrawal 2012). We run 10 000 bootstrap rounds for each replicate line. In each round, we (1) drew three bootstrap samples *(i.e.* sampling with replacement, with sample size equal to the size of the sampled set): one from the set of control sires (26, 19 and 14 sires for lines 50, 52, and 55, respectively), one from the set of 20 Gy-irradiated sires (28, 19, and 14 sires, respectively), and one from the set of 40 Gy-irradiated sires (26, 20 and 13 sires, respectively); (2) noted fitness scores of son and daughter of each sampled sire; (3) calculated mean fitness separately for each dose and sex, based on the bootstrap samples; and (4) calculated sm, sf, and their ratio (see Eq. 1 and below) separately for 20 and 40 Gy. For each of the metrics of interest (S_M_, S_F_, and S_M_/S_F_), the 2.5^th^ and 97.5^th^ percentiles of the 10 000 bootstrap scores were then taken as the lower and the upper limit, respectively, of the confidence interval.

### Estimating sexual selection against induced mutations

Barring any catastrophes, the following things were happening in each of the experimental “populations” in the male fitness assay: females produced eggs, fertilizations of these eggs were shared between a focal male and his competitors, and a certain fraction of the fertilized eggs survived to adulthood, at which stage we determined their paternity (focal *vs*. rival), and scored the focal male’s fitness as the number of his offspring. Thus, detrimental effects of mutations on male fitness could come about at either or both of the two stages: during competition with rival males for eggs (sexual selection acting on the focal males) and/or during offspring development from fertilized egg to imago (viability selection acting on the males’ offspring).

We estimated the specific contribution of sexual selection, separately for each replicate line, and only for the 40 Gy treatment (because in the 20 Gy group we did not detect any significant decline in male fitness). We calculated the (unstandardized) coefficients of sexual selection (S_s_) as differences in mean number of *rival* males’ offspring between the mutated and control treatments (cf. Appendix, eq. B2). In Appendix we explain the full rationale for this approach; briefly: the number of rivals’ offspring is affected by the focal males’ sexual competitiveness (the less competitive the focal males, the more eggs get fertilized by their rivals and *vice versa*) but not by the viability of the focals’ offspring. Thus, the difference in mean number of rival males’ offspring between the mutated and control treatments reflects the strength of sexual selection against the induced mutations, unconfounded by the effect of selection acting at the level of offspring viability. We used t tests to assess if the effects of sexual selection were significantly different from 0, which would indicate that the induced mutations significantly decrease male sexual competitiveness. To assess the contribution of sexual selection relative to the overall selection against the induced mutations (as measured in our assays), we also calculated the unstandardized coefficients of total selection (S_t_) as differences between the mean number of focal males’ offspring between the control and mutated treatments (cf. Appendix, eq. B2). We then calculated the ratio of sexual selection to total selection coefficients (S_s_/S_t_). We first calculated Ss, St, and their ratio separately for each replicate line. Subsequently, to obtain an overall estimate for all replicates, we used ANOVAs to analyze the effects of mutation treatment (40 Gy vs. control), replicate line (50, 52 and 55), and their interaction on the number of (1) rival and (2) focal offspring. We then calculated Ss and St based on least squares means for irradiation treatment groups, extracted from model (1) and (2), respectively.

### Behavioural assay

The assay was carried out in the same CT room in which all our beetles are normally housed, in order to minimize the impact of the assay conditions on normal behavior of the beetles. Each experimental male (from either 40 Gy or control treatment) was placed on an observation arena along with one virgin female from the stock culture and one virgin RdHD male. To differentiate between experimental males and females during the behavioral assay, each experimental male was marked with a tiny drop of white correction fluid on the thorax (RdHD males are easily distinguished by their antennal morphology). Observation arenas were 6 cm Petri dishes filled with small amount of culture medium (flour:yeast mixture and rolled oats)-the amount was adjusted so that the beetles could feed and move around without slipping on plastic, while remaining visible at all times. Each such group of beetles was observed for 1 hour, during which the experimental male’s behavior was scored every minute as either 1 (mounting the female) or 0 (not mounting). From these scores, for each male we obtained two measures of sexual activity: the latency to first mounting, and the total number of times it was observed mounting a female (which would be closely related to the total time spent mounting).

Once again, we analyzed the data separately for each replicate line. Because of very skewed distributions, we used Mann-Whitney tests to compare latency to first mounting and total mounting time between the control and mutated males. For lines 52 and 55, the variance of total mounting time differed significantly between treatments; therefore, we took logarithm of the data before the analysis.

## Results

### Selection against induced mutations in males and females

The 20 Gy dose caused significant decline in female fitness in two lines (50 and 52), but no effect in the third, whereas male fitness showed no significant decrease in any of the lines (Fig. 1, top row). The 40 Gy dose caused significant decline in fitness of both sexes, in all 3 lines (Fig. 1, bottom row). The coefficients of selection against mutations introduced by 40 Gy dose were higher in females in all lines, resulting in m/f ratios smaller than 1, but the trend was not significant in any line, and the confidence intervals around m/f were very wide in two lines (Fig. 1, bottom row). The same trend (m/f<1) was observed under 20 Gy dose in lines 50 and 52.

### Selection against induced mutations in males: the role of sexual selection

Sexual selection coefficient was positive for all lines, but not significantly greater than 0 in line 50 (S_s_ = 10.04; t_50_ = 1 27, P = 0.21, S_s_/S_t_ = 0.28) nor in line 55 (when including all data: S_s_ = 1.63; t_23_ = 0.10, P = 0.92, S_s_/S_t_ = 0.04; after excluding an influential outlier: S_s_ = 13.91; t_22_ = 1.34, P = 0.19, S_s_/S_t_ = 0.30). In line 52, sexual selection coefficient was significant (S_s_ = 28.86; t_32_ = 4.50, P<0.001) and, in fact, larger than the total selection coefficient (S,_s_/S_t_ = 1.36). When analyzing data from all lines together using ANOVA, the effect of sexual selection against induced mutations (estimated as the effect of mutation treatment on the number of rival offspring, see Methods) was significant (when including all data, mutation treatment: F_1,105_ = 6.47, P = 0.012, replicate line: F_2,105_ = 2.38, P = 0.098, interaction: F_2,105_ = 1.79, P = 0.172; after excluding an influential outlier, F_1,104_ = 12.55, P = 0.001, replicate line: F_2,104_ = 1.72, P = 0.184, interaction: F_2,104_ = 1.49, P = 0.229). The coefficient of sexual selection (calculated based on ANOVA least squares means for mutation treatment groups) was S_s_ = 13.51 (S_s_/S_t_ = 0.43) when all data was included and S_s_ = 17.61 (S_s_/S_t_ = 0.51) if an influential outlier was excluded.

**Figure 1.**
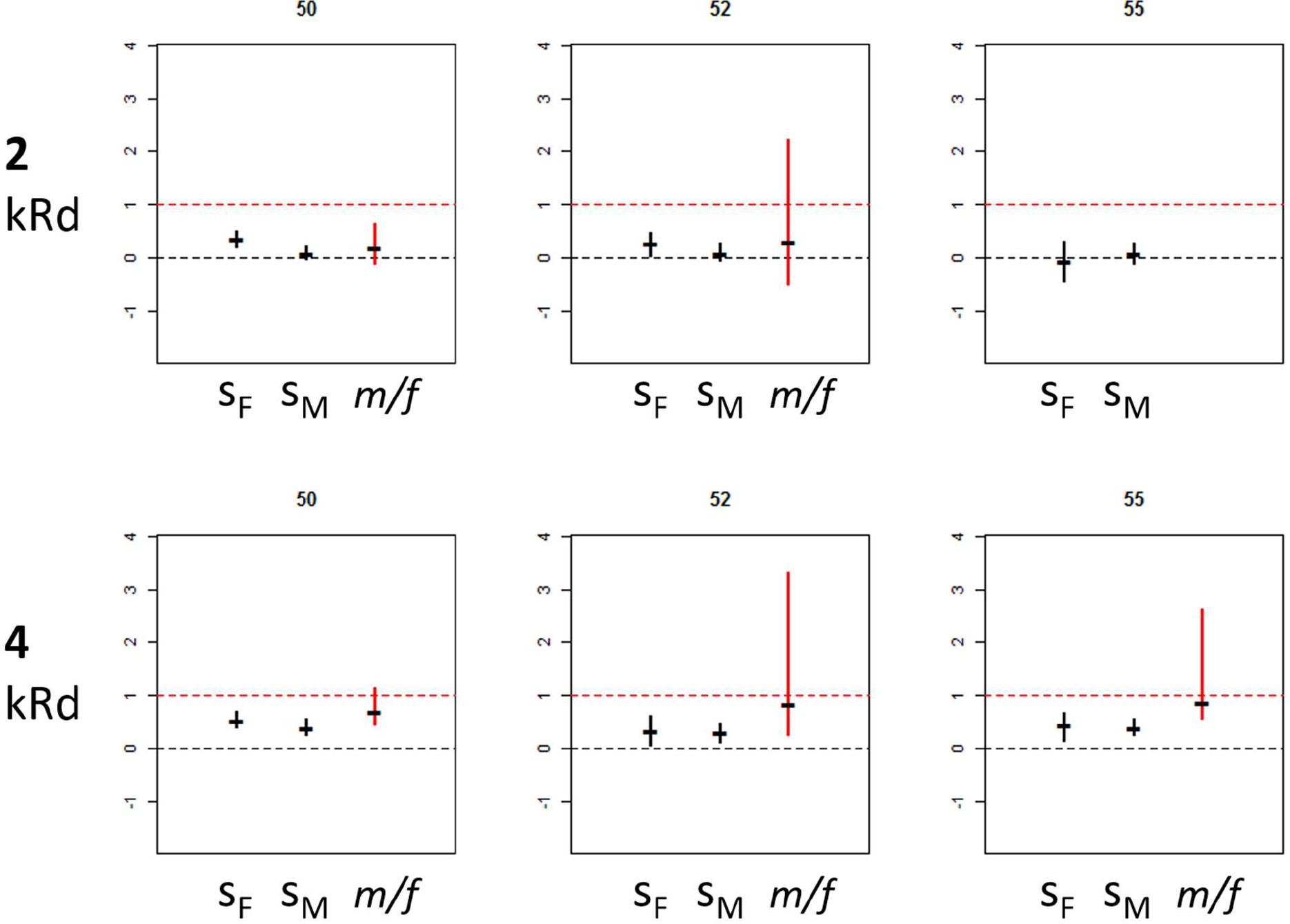
Standardized coefficients of selection against induced mutations in females (S_F_) and males (S_M_), and the ratio S_M_/S_F_ (m/f), ± bootstrap confidence intervals, in the 20 Gy (top row) and 40 Gy (bottom row) treatments.

In the behavioral assay, we found no difference between the control and mutated treatments in either line in mean latency to the first mount (line 50: W = 244, P = 0.689; line 52: W = 118.5, P = 0.968; line 55: W = 64, P = 0.742), nor in the mean time spent mounting (line 50: W = 247.5, P = 0.746; line 52: W = 116, P = 0.984; line 55: W = 73, P = 0.883). In lines 52 and 55, the variances of total mounting time were significantly higher in the mutated treatment (line 52: F = 6.817, P = 0.0004; line 55: F = 10.62, P = 0.0002).

## Discussion

Sexual selection on males can contribute to the maintenance of sex by reducing deleterious mutation load in sexual populations relative to their asexual competitors, provided that it discriminates against the same alleles as selection acting on females, and does so with higher efficiency (Manning 1984; Agrawal 2001; Siller 2001; Whitlock and Agrawal 2009). Our results suggest that the latter condition is not satisfied in *T. castaneum*.

We studied selection against deleterious mutations introduced by ionizing radiation and present in the genome in heterozygous state (hence mimicking the natural situation whereby individual deleterious alleles are rare), in the context of small experimental populations with 1:1 sex ratio (mimicking the sex ratio of natural populations). In the 20 Gy treatment, we found no effect of induced mutations on male fitness in any replicate, whereas female fitness declined significantly in two replicate lines. In the 40 Gy treatment, both female and male fitness declined in all three replicates; however, there was no evidence of more detrimental effect of mutations on males. In fact, the trend was in the opposite direction (female fitness being affected more strongly in all three lines), although it was non-significant. Deleterious mutations on the X chromosome could only have negligible contribution to this result, since X only constitutes *ca*. 6% of *T. castaneum* genome (Trauner et al. 2009).

When estimated across replicate lines, the effect of sexual selection constituted 43% or 51% (depending on the inclusion of an influential outlier) of overall selection against induced mutations. The within-line estimates were all positive, meaning that sexual competitiveness of mutated males was consistently lower across the three replicate lines (further supported by the lack of mutation × line interaction effect in the ANOVA), although the magnitude of Ss varied among the lines and in two of them it was not significantly different from zero. In the third line, the contribution of sexual selection was significant, and, surprisingly, exceeded the estimate of overall selection, indicating that egg-to-adult survival of the focal males’ offspring was higher in the mutated than in the control treatment. The measures of male sexual activity did not differ in means between the control and mutated treatment in either line (although in two lines, the variances in total time spent in copulatory mounts were significantly higher in the mutated treatment). Thus, we speculate that the observed effect of sexual selection against induced mutations may be due to decreased sperm competitiveness of the mutated males. In line with this speculation, sperm competitiveness in *T. castaneum* was previously shown to be affected by mutations revealed by inbreeding (Michalczyk et al. 2010).

Taken together, our results indicate that sexual selection on males does discriminate against mutations that are also detrimental to female fitness, although we were not able to pinpoint the exact mechanism involved. However, we show that sexual selection does not make these mutations more deleterious in males than in females, indicating that it does not contribute to offsetting the costs of sex.

These findings contrast with some of the conclusions reached in a recent study on the same species by Lumley et al. (2015). They created two experimental evolution regimes within which they exposed replicate populations to either decreased (monogamy or female-biased sex ratio) or elevated (polyandry or male-biased sex ratio) levels of sexual selection. Subsequently, they exposed the load of recessive and partly recessive mutations present in the evolved populations by creating inbred lines (20 generations of sib × sib mating). They found that inbred lines derived from populations with the history of strong sexual selection showed slower fitness decline over generations, and survived longer, than the lines derived from populations with weak/eliminated sexual selection, indicating that mutation load in traits affecting survival and offspring production was higher in the latter. This provides compelling evidence that the net directions of sexual and “ordinary” (Darwin 1859) selection are aligned, rather than antagonistic, which has important theoretical as well as potential practical implications (Rowe and Houle 1996; Tomkins et al. 2004; Holman and Kokko 2013; Charge et al. 2014). However, it is not enough to “provide compelling support for the (…) models (…) which argued that costs of sex could be offset by population genetic benefits derived from sexual selection” (Lumley et al. 2015). This is because relaxing sexual selection on males will hamper purging mutation load in female and population fitness (thus creating the difference in load level between populations differing in the strength of sexual selection) whenever the net effect of selection on genome is in the same direction for males and females, regardless of its relative strength between the sexes (Whitlock and Agrawal 2009). Yet, selection being stronger in males is a key condition in the theoretical models featuring sexual selection as a contributor to the maintenance of sex (Agrawal 2001; Siller 2001; Whitlock and Agrawal 2009 cf. Introduction). Our results suggest, instead, that mutation load in *T. castaneum* is similar, or even higher, than it would be under a (hypothetical) scenario of asexual reproduction (Whitlock and Agrawal 2009, see also Introduction). The results obtained by Lumley et al. (2015) could thus be due to the enhanced SS populations reducing the mutation load below-and/or the relaxed SS populations accumulating the load above-this level.

That being said, we need to acknowledge two main limitations of our experimental approach. First, we analyzed the effects of mutations artificially induced by ionizing radiation, which may have different effects from these occurring naturally. However, ionizing radiation induces mutations with a wide range of effects (e.g. Evans and DeMarini 1999), and the distribution of these effects is not likely to be fundamentally different from that characterizing spontaneous mutations (Radwan 2004). Hence, there is no *a priori* reason to expect that induced mutations will substantially differ from spontaneous ones in their relative effects on male *vs.* female fitness. Second, as a background to assess the reproductive success of our experimental males and females we used the reindeer (RdHD) strain. This allowed us to distinguish wild type and RdHD offspring, and hence score the reproductive success of our focal beetles while letting them interact with other individuals at the 1:1 sex ratio-the motivation being to mimic, as closely as logistically possible, the conditions normally experienced in populations. Competition assays between these two strains (wild type and RdHD) are commonly used while working with flour beetles to assess the fitness of experimental wild type males or examine paternity shares after multiple copulations (Lewis et al. 2005; Michalczyk et al. 2011; Demont et al. 2014). However, RdHD males are relatively poor competitors when faced with wild type males (in this study, control males sired on average over 50% offspring, and mutated males-*ca.* 40% offspring, in competition with 3 RdHD rivals). In other words, the RdHD males provided a mild competitive environment for the focal males, which may have resulted in masking the negative effects of induced mutations and hence-in underestimating the selection coefficients for males. However, although it is counterintuitive, selection needs not necessarily be weaker in milder environments (Agrawal and Whitlock 2010). Nevertheless, an ideal fitness assay would examine competition between both mutated and control wild type males. This could be achieved in future studies by inducing mutations into an outbred population, using (non-mutated) individuals from the same population as rivals in fitness assays, and then applying molecular tools to assign offspring parentage to focal *vs.* rival individuals.

In summary, we would like to emphasize the importance of comparing male and female selection coefficients when studying the role of sexual selection in the evolution of sex. To date, such comparisons are relatively rare (see Introduction), despite being central to testing the relevant models (Agrawal 2001; Siller 2001).

## Appendix

We made two simplifying assumptions: (1) that the total number of fertilized eggs in a “population” is unaffected by focal male’s treatment and (2) that survival to adulthood of eggs fertilized by control focal males and rival males is the same. The first assumption seems reasonable, as the total number of fertilized eggs in our experimental “populations” is likely determined by the number of eggs produced by females (given anizogamy and 1:1 sex ratio). The second assumption is likely to be met in our study, as there appear to be no difference in egg-to-adult survival between the wild type and Rd strains (Michalczyk, unpublished results, Michalczyk et al. 2010). Based on these assumptions, we can parametrize the male fitness assay’s dynamics as follows:

| Variable | Control treatment | Mutated treatment |
| --- | --- | --- |
| total # of fertilized eggs in a “population” | a | a |
| % of *a* fertilized by a focal male | c | d |
| % of *a* fertilized by rival males | 1-c | 1-d |
| % surviving to adulthood out of all eggs fertilized by a focal male | e | f |
| % surviving to adulthood out of all eggs fertilized by rival males | e | e |
| from the above, we can calculate the following: |  |  |
| # eggs fertilized by a focal male ($Ef$) | ac | ad |
| # eggs fertilized by rival males ($Er$) | a(1-c) | a(1-d) |
| # adult offspring of a focal male ($Of^*$) | ace | adf |
| # adult offspring of rival males ($Or$) | a(1-c)e | a(1-d)e |
| total # adult offspring in a “population” ($Ot$) | ace + a(1-c)e<br>=<br>ae | adf + a(1-d)e = adf<br>+ ae - ade |

If we further assume that the variables a-f are all independent of each other, then the expected (mean) values of the last five variables are

| Variable mean | Control treatment | Mutated treatment |
| --- | --- | --- |
| $\overline{Ef}$ | $\bar{a} \bar{c}$ | $\bar{a} \bar{d}$ |
| $\overline{Er}$ | $\bar{a} (1 - \bar{c})$ | $\bar{a} (1 - \bar{d})$ |
| $\overline{Of^*}$ | $\bar{a} \bar{c} \bar{e}$ | $\bar{a} \bar{d} \bar{f}$ |
| $\overline{Or}$ | $\bar{a} (1 - \bar{c}) \bar{e}$ | $\bar{a} (1 - \bar{d}) \bar{e}$ |
| $\overline{Ot}$ | $\bar{a} \bar{e}$ | $\bar{a} \bar{d} \bar{f} + \bar{a} \bar{e} - \bar{a} \bar{d} \bar{e}$ |

Sexual selection against the induced mutations is manifested as the difference between the control and mutated focal males in the mean fraction 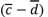 and, in consequence, in the mean number 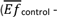 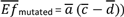 of fertilized eggs. Sadly, these variables cannot be measured in our experiment (nor in many other experiments following similar fitness assay design): only *Of, Or* and their sum (*Ot*) can actually be scored in the assay.
At the stage of adult offspring, selection is measured as:

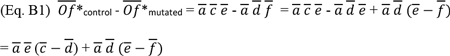

The first part of the equation 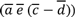 can be considered a proper measure of sexual selection against the induced mutations, albeit scored at the stage of adult offspring rather than fertilized eggs. This is because it arises from the difference in fertilization success between the control and mutated males, scaled only by two quantities which are independent of the mutation treatment (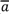 and 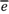). In other words: mutation treatment influences this quantity only via the effects on fertilization success, whereas the total difference between 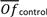 and 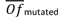 is additionally influenced by the mutations’ effect on offspring survival.

We can estimate 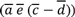 by calculating the difference in the number of rival-rather than focal - offspring between the mutated and control treatments:

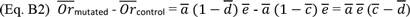

(Note that the *control* and *mutated* subscripts in Eq. B2 refer to treatment groups and not to rival males themselves, which are not mutated in any treatment group).

Finally, the second part of Eq. B1 is equal to the difference between the control and mutated treatments in the mean total number of adult offspring produced by the experimental “populations””

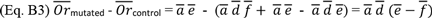

This reflects the fact that decreased survival of the mutated focal males’ offspring leads to a decline in the total number of adult offspring produced by experimental “populations” in the mutated treatment (if offspring survival is unaffected by treatment, then 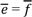 and Eq. B3 gives 0: there is no difference between treatments in the total offspring number produced by “populations”).

* Of here is equivalent to W in the main text, because we take the number of adult offspring as a measure of the focal males’ fitness

## Acknowledgments

We are grateful to Lukasz Michalczyk, Wieslaw Babik and the members of Molecular and Behavioural Ecology Group (Jagiellonian University) for their comments on the manuscript, and to Michael Morrissey and Magdalena Herdegen for discussion on (some of) the analyses. This work was supported by the Ministry of Science and Higher Education grant IP2012 044872 to Z.M.P. and by the Jagiellonian University grant DS/WBiNoZ/INoS/762/15. We declare no conflict of interests.

## References

Agrawal, A. F. 2001. Sexual selection and the maintenance of sexual reproduction. Nature 411:692–695

Agrawal, A. F., and M. C. Whitlock. 2010. Environmental duress and epistasis: how does stress affect the strength of selection on new mutations? Trends Ecol. Evol. 25:450–458

Almbro, M., and L. W. Simmons. 2013. Sexual selection can remove an experimentally induced mutation load. Evolution 68:295–300

Andersson, M. 1994. Sexual selection. Princeton University Press, Princeton

Charge, R., C. Teplitsky, G. Sorci, and M. Low. 2014. Can sexual selection theory inform genetic management of captive populations? A review. Evol. Appl. 7:1120–1133

Connallon, T., and A. G. Clark. 2014. Evolutionary inevitability of sexual antagonism. Proc. R. Soc. B Biol. Sci. 281:20132123

Darwin, C. 1859. On the origin of species. John Murray, London

Demont, M., V. M. Grazer, L. Michalczyk, A. L. Millard, S. H. Sbilordo, B. C. Emerson, M. J. G. Gage, and O. Y. Martin. 2014. Experimental removal of sexual selection reveals adaptations to polyandry in both sexes. Evol. Biol. 41:62–70

Duffy, E., R. Joag, J. Radwan, N. Wedell, and D. J. Hosken. 2014. Inbreeding alters intersexual fitness correlations in Drosophila simulans. Ecol. Evol. 4:3330–3338

Evans, H. H., and D. M. DeMarini. 1999. Ionizing radiation-induced mutagenesis: radiation studies in Neurospora predictive for results in mammalian cells. Mutat. Res. 437:135–150

Grieshop, K., J. Stångberg, I. Martinossi-Allibert, G. Arnqvist, and D. Berger. 2016. Strong sexual selection in males against a mutation load that reduces offspring production in seed beetles. J. Evol. Biol.doi: 10.1111/jeb.12862.

Hetfield, J., L. Ulrich, and D. Mustaine. 1983. Seek & Destroy (in: Kill‘Em All). Music America Studios in Rochester. New York.

Hollis, B., and D. Houle. 2011. Populations with elevated mutation load do not benefit from the operation of sexual selection. J. Evol. Biol. 24:1918–1926

Holman, L., and H. Kokko. 2013. The consequences of polyandry for population viability, extinction risk and conservation. Philos. Trans. R. Soc. B Biol. Sci. 368:201230053

Keightley, P. D., and M. Lynch. 2003. Toward a realistic model of mutations affecting fitness. Evolution 57:683–685

Kotiaho, J. S., and M. Puurtinen. 2007. Mate choice for indirect genetic benefits: scrutiny of the current paradigm. Funct. Ecol. 21:638–644

Laskowski, R., J. Radwan, K. Kuduk, M. Mendrok, and P. Kramarz. 2015. Population growth rate and genetic variability of small and large populations of red flour beetle (Tribolium castaneum) following multigenerational exposure to copper. Ecotoxicology 24:1162–1170

Lewis, S. M., A. Kobel, T. Fedina, and R. W. Beeman. 2005. Sperm stratification and paternity success in red flour beetles. Physiol. Entomol. 30:303–307

Lewis, S. M., N. Tigreros, T. Fedina, and Q. L. Ming. 2012. Genetic and nutritional effects on male traits and reproductive performance in Tribolium flour beetles. J. Evol. Biol. 25:438–451

Lumley, A. J., Ł. Michalczyk, J. J. N. Kitson, L. G. Spurgin, C. A. Morrison, J. L. Godwin, M. E. Dickinson, O. Y. Martin, B. C. Emerson, T. Chapman, and M. J. G. Gage. 2015. Sexual selection protects against extinction. Nature 522:470–473

Mallet, M. A., and A. K. Chippindale. 2011. Inbreeding reveals stronger net selection on Drosophila melanogaster males: implications for mutation load and the fitness of sexual females. Heredity 106:994–1002

Manning, J. T. 1984. Males and the advantage of sex. J. Theor. Biol. 108:215–220

McGuigan, K., D. Petfield, and M. W. Blows. 2011. Reducing mutation load through sexual selection on males. Evolution 65:2816–2829

Michalczyk, Ł., O. Y. Martin, A. L. Millard, B. C. Emerson, and M. J. G. Gage. 2010. Inbreeding depresses sperm competitiveness, but not fertilization or mating success in male Tribolium castaneum. Proc. R. Soc. B Biol. Sci. 277:3483–3491

Michalczyk, Ł., A. L. Millard, O. Y. Martin, A. J. Lumley, B. C. Emerson, and M. J. G. Gage. 2011. Experimental evolution exposes female and male responses to sexual selection and conflict in Tribolium castaneum. Evolution 65:713–724

Milinski, M. 2006. The major histocompatibility sexual selection, complex, and mate choice. Annu. Rev. Ecol. Evol. Syst. 37:159–186

Pekkala, N., M. Puurtinen, and J. S. Kotiaho. 2009. Sexual selection for genetic quality: disentangling the roles of male female behaviour and female behaviour. Anim. Behav. 78:1357–1363

Pischedda, A., and A. K. Chippindale. 2006. Intralocus sexual conflict diminishes the benefits of sexual selection. PLoS Biol. 4:2099–2103

Plesnar, A., M. Konior, and J. Radwan. 2011. The role of sexual selection in purging the genome of induced mutations in the bulb mite (Rizoglyphus robini). Evol. Ecol. Res. 13:209–216

Plesnar-Bielak, A., A. M. S krzynecka, K. Miler, and J. Radwan. 2014. Selection for alternative male reproductive tactics alters intralocus sexual conflict. Evolution 68:2137–2144

Power, D. J., and L. Holman. 2015. Assessing the alignment of sexual and natural selection using radiomutagenized seed beetles. J. Evol. Biol. 28:1039–1048

Price, T., D. Schluter, and N. E. Heckman. 1993. Sexual selection when the female directly benefits. Biol. J. Linn. Soc. 48:187–211

Radwan, J. 2004. Effectiveness of sexual selection in removing mutations induced with ionizing radiation. Ecol. Lett. 7:1149–1154

Radwan, J., J. Unrug, K. Śnigórska, and K. Gawrońska. 2004. Effectiveness of sexual selection in preventing fitness deterioration in bulb mite populations under relaxed natural selection. J. Evol. Biol. 17:94–99

Rowe, L., and D. Houle. 1996. The lek paradox and the capture of genetic variance by condition dependent traits. Proc. R. Soc. B Biol. Sci. 263:1415–1421

Sharp, N. P., and A. F. Agrawal. 2012. Male-biased fitness effects of spontaneous mutations in Drosophila melanogaster. Evolution 67:1189–1195

Sharp, N. P., and A. F. Agrawal. 2008. Mating density and the strength of sexual selection against deleterious alleles in Drosophila melanogaster. Evolution 62:857–867

Shuker, D. M. 2010. Sexual selection: endless forms or tangled bank? Anim. Behav. 79:e11–e17

Siller, S. 2001. Sexual selection and the maintenance of sex. Nature 411:689–692

Tomkins, J. L., J. Radwan, J. S. Kotiaho, and T. Tregenza. 2004. Genic capture and resolving the lek paradox. Trends Ecol. Evol. 19:323–328

Trauner, J., J. Schinko, M. D. Lorenzen, T. D. Shippy, E. A. Wimmer, R. W. Beeman, M. Klingler, G. Bucher, and S. J. Brown. 2009. Large-scale insertional mutagenesis of a coleopteran stored grain pest, the red flour beetle Tribolium castaneum, identifies embryonic lethal mutations and enhancer traps. BMC Biol. 7:1–12

Whitlock, M. C., and A. F. Agrawal. 2009. Purging the genome with sexual selection: reducing mutation load through selection on males. Evolution 63:569–582

Whitlock, M. C., and D. Bourguet. 2000. Factors affecting the genetic load in Drosophila: synergistic epistasis and correlations among fitness components. Evolution 54:1654–1660

